# Dose-dependent effects of netarsudil, a Rho-kinase inhibitor, on the distal outflow tract

**DOI:** 10.1101/2020.01.17.909101

**Authors:** Si Chen, Susannah Waxman, Chao Wang, Sarah Atta, Ralitsa Loewen, Nils A. Loewen

**Affiliations:** University of Pittsburgh School of Medicine, Department of Ophthalmology, Pittsburgh, Pennsylvania, United States; Xiangya Hospital of Central South University, Department of Ophthalmology, Changsha, Hunan, China; University of Würzburg, Department of Ophthalmology, Würzburg, Germany

**Author notes:** Corresponding Author: Nils A. Loewen, MD, PhD, Department of Ophthalmology, University of Würzburg, Josef-Schneider-Straße 11, 97080 Würzburg, Germany. **Grant information:** National Eye Institute K08EY022737 (NAL); Initiative to Cure Glaucoma of the Eye and Ear Foundation of Pittsburgh (NAL); P30-EY08098 (NAL); Department grant by Research to Prevent Blindness (NAL); an unrestricted fellowship grant from the Xiangya Hospital of Central South University (SC).

**Keywords:** Rho-kinase inhibitor, netarsudil, distal outflow tract, anterior chamber perfusion model, porcine eyes

## Abstract

**Purpose:** To characterize the effects of netarsudil on the aqueous humor outflow tract distal to the trabecular meshwork (TM). We hypothesized that netarsudil increases outflow facility in eyes with and without circumferential ab interno trabeculectomy (AIT) that removes the TM.

**Methods:** 64 porcine anterior segment cultures were randomly assigned to groups with (n=32) and without circumferential AIT (n=32). Cultures were exposed to 0.1, 1, and 10 μM netarsudil (N= 8 eyes per concentration). For each concentration, IOP and vessel diameters were compared to their respective pretreatment baselines. Outflow tract vessel diameters were assessed by spectral-domain optical coherence tomography (SDOCT) and rendered in 4D (XYZ time-series).

**Results:** Netarsudil at 1 μM reduced IOP in both eyes with TM (−0.60±0.24 mmHg, p = 0.01) and in eyes without TM (−1.79±0.42 mmHg, p<0.01). At this concentration, vessels of the distal outflow tract dilated by 72%. However, at 0.1 μM netarsudil elevated IOP in eyes with TM (1.59±0.36 mmHg, p<0.001) as well as in eyes without TM (0.23±0.32 mmHg, p<0.001). Vessels of the distal outflow tract constricted by 31%. Similarly, netarsudil at a concentration of 10 μM elevated IOP both in eyes with TM (1.91±0.193, p<0.001) and in eyes without TM (3.65±0.86 mmHg, p<0.001). At this concentration, outflow tract vessels constricted by 27%.

**Conclusion:** In the porcine anterior segment culture, the dose-dependent IOP changes caused by netarsudil matched the diameter changes of distal outflow tract vessels. Hyper- and hypotensive properties of netarsudil persisted after TM removal.

## Introduction

Primary open-angle glaucoma (POAG) is a leading cause of irreversible blindness, with 42% of patients eventually losing vision in one eye [1]. The annual costs of glaucoma in the US are $5.8 billion [2]. Most POAG is treated with eye drops, but even the latest prostaglandin analogs offer continuous-treatment success rates of 10% at one year [3]. Increased, optic nerve-damaging intraocular pressure (IOP) in POAG was long thought to be only caused by outflow resistance at the trabecular meshwork (TM), which guards the drainage system of the eye. However, data from clinical TM ablation in thousands of patients show it fails to lower IOP to the pressure level in the recipient episcleral veins [4–8]. The data suggests that over half of resistance resides in the distal outflow tract (OT), downstream of the TM and Schlemm’s canal (SC). The loci and substrates of such distal outflow resistance are unknown, but critical to identify. New evidence of post-TM outflow regulatory structures was recently presented, using automatic 3D segmentation and outflow reconstruction with wide-spectrum spectral-domain optical coherence tomography (SD-OCT). Outflow vessel dilation by nitric oxide correlated to a 61.5% increased outflow in porcine [9] and human eyes [10]. Recent studies of intracameral bimatoprost suggest similar effects [11].

Present ocular hypotensive medications either reduce aqueous humor production (beta-blockers, alpha-agonist, or carbonic anhydrase inhibitors) or increase the uveoscleral outflow as the main mechanism (prostaglandin analogs) [12]. Older muscarinic substances like pilocarpine have a direct effect on trabecular flow but fell out of favor because of their side effects that include pupillary constriction and myopization of phakic patients [13].

In this study, we examined netarsudil, an inhibitor of Rho-kinase, and the norepinephrine transporter [13]. It can increase the outflow facility by expanding the juxtacanalicular TM and by dilating the episcleral veins [14, 15]. It was approved by the US Food and Drug Administration in December 2017 as a 0.02% daily single-dose medication and is currently in phase 3 studies in Europe [16]. Contradictory observations in pilot experiments made us hypothesize that there are dose-dependent effects on the distal outflow tract that could be discovered in ex vivo porcine anterior segment cultures after removing the trabecular meshwork.

## Materials and Methods

### Study Design

We studied the outflow facility responses of anterior segment organ cultures to 0.1, 1, and 10 μM netarsudil. To establish the contribution by the TM to these responses, treatment groups were created for each concentration with and without circumferential ab interno trabeculectomy (AIT) using a trabectome as described before [17]. Eight eyes were randomly assigned to each treatment group and perfused for at least 48 hours to establish a stable baseline IOP before treatment.

We determined the structural response of outflow tract vessels distal to the TM at the same concentrations of 1 of 0.1, 1, and 10 μM netarsudil using two eyes per treatment group. Time-series volumetric scans of the perilimbal region were captured via wide-spectrum spectral-domain optical coherence tomography (SD-OCT) pre and post-treatment. Outflow tract cross-sectional areas were compared to respective baselines.

### Materials

Porcine eyes were obtained from a local abattoir (Thoma Meat Market, Saxonburg, Pittsburgh PA) and cultured within two hours of sacrifice. Extraocular tissues, including the conjunctiva, were carefully removed. Eyes were decontaminated by submersion in 5% povidone-iodine ophthalmic solution (Betadine 5%, Fisher Scientific, NC9771653) for two minutes, and hemisected in a biosafety cabinet. After removal of the posterior segment, lens, and ciliary body, anterior segments were mounted on custom perfusion dishes.

### Ocular perfusion and outflow measurement

Anterior segments were cultured at 37° C and perfused at 4 μl/min with media (Dulbecco’s modified Eagle medium (DMEM); sh30284.02, Fisher Scientific), 1% FBS (10082-147; Fisher Scientific), 1% antibiotic, antimycotic (15240-062; Fisher Scientific), with a microinfusion pump (70-3007; Harvard Apparatus, Holliston, MA, USA). IOP was measured at 2-minute intervals with pressure transducers (Deltran II: DPT-200; Utah Medical Products, Midvale, UT, USA), and recorded (FE224, PL3508/P, MLA1052; ADInstruments, Sydney, Australia; LabChart 7; ADInstruments) [17]. Baseline IOP was achieved after 48 hours of perfusion. The effect of netarsudil on IOP was observed over the subsequent time.

### SD-OCT imaging and analysis

Each porcine eye was positioned with the optic nerve remnant secured in a low-compression mount (CryoElite Cryogenic Vial #W985100; Wheaton Science Products, Millville, NJ, USA) and kept damp with phosphate-buffered saline. Anterior chambers were perfused at a constant pressure of 15 mmHg with perfusion media as done previously [17]. The eyes were placed under the sample arm of an SD-OCT equipped with a 10 mm telecentric lens (Envisu R2210, Leica, Bioptigen, Morrisville, NC, USA). The scanning beam was oriented perpendicularly to a portion of the limbus in which intrascleral signal voids of the outflow tract could be visualized. After 30 minutes of perfusion to stabilize outflow, volumetric baseline scans (6×4×1.6 mm) were captured. The medium was then supplemented with 0.1, 1, or 10 μM netarsudil, and a gravity-mediated anterior chamber exchange was performed. Two eyes were imaged for each treatment group in a single session. To minimize imaging artifacts and registration errors, adjustment of the SD-OCT sample arm was limited. This permitted scans at 30-minute increments in one eye for each treatment group without readjusting the sample arm. The other eye was imaged only at baseline and at the end of the 3-hour experiment after careful sample arm readjustment. Each scan created 600 images, resulting in at least 5,400 images analyzed per group (0.1, 1, and 10 μM netarsudil).

### Trabecular meshwork removal

The TM was removed by a glaucoma surgeon experienced in ab interno trabeculectomy. Anterior segments were placed under an ophthalmic surgery microscope (S4, Carl Zeiss Meditec, Jena, Germany) and positioned with the cornea facing down in an aseptic holder. TM removal was then performed over the entire circumference via Trabectome (Neomedix Corp, Tustin, California, USA) as described before [4]. TM removal was confirmed by histology.

### Data Analysis

IOP measurements were down-sampled into 2-hour blocks and normalized to respective controls. Pre-treatment was compared to post-treatment with a one-sample t-test in Python 3.6 [18]. SD-OCT images were processed in ImageJ [19] (Version 1.50i, National Institute of Health, Bethesda, Maryland, United States) and Amira Aviso (version 9.1, FEI, ThermoScientific, Waltham, Massachusetts) as done previously [17] to remove noise, align pre-and post-treatment outflow tract signal voids in a three-dimensional space (3D), and to allow automated, quantitative measurement of cross-sectional areas (CSA). Cross-section areas of pre- and post-treatment were compared with a student’s t-test.

## Results

Histology of eyes in which the porcine TM was left intact presented with the characteristic appearance of the aqueous angular plexus and several circumferential canal elements (**Figure 1 A**). The TM appeared as a prominent, multilayered structure with trabecular beams populated by TM cells, which became more condensed towards the canal elements. Histology of eyes that had passed through the experiment and in which the TM had been ablated by AIT lacked the TM (**Figure 1 B**). Circumferentially running, sagittally cut, canal-like elements could be seen adjacent to the space where the TM had been removed.

**Figure 1:**
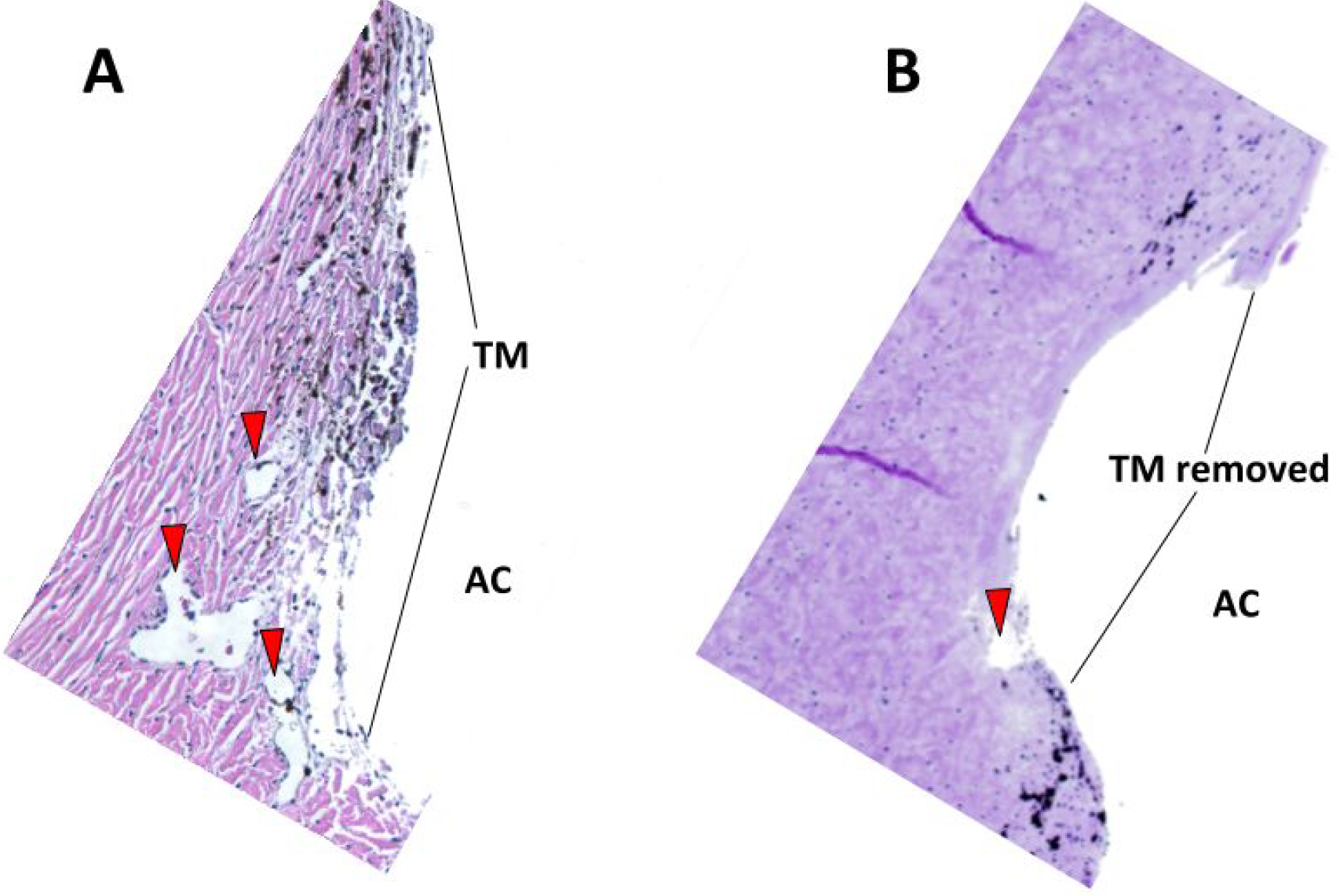
Histology of the porcine angular aqueous plexus of perfusion cultured anterior segments rotated to surgeons view. A) A section from a non-ablated eye shows an intact trabecular meshwork (TM) and sagittally cut, circular drainage channel segments (red arrows). B) The section from an eye with circumferentially ablated TM by AIT obtained after perfusion experiments with netarsudil. The TM is removed and circumferential drainage channels are partially unroofed.

Eyes with an intact TM that were exposed to 0.1 μM netarsudil experienced an IOP elevation by 1.59±0.36 mmHg (p < 0.001, **Figure 2**) when compared to baseline. In contrast, eyes perfused with netarsudil at a concentration of 1 μM experienced a reduction of IOP by −0.60±0.24 mmHg (80.31±48.21% reduction, p = 0.01) reduction. We observed again a significant IOP elevation of 1.91±0.19 (p < 0.001) at a higher concentration of 10 μM. Eyes that had undergone a circumferential removal of TM by AIT showed an IOP elevation by 0.23±0.32 mmHg at 0.1 μM netarsudil (p<0.001, **Figure 2**), just like eyes with an intact TM. However, IOP was lowered by 1 μM netarsudil (−1.79±0.42 mmHg, p < 0.001), as seen in non-ablated eyes. At 10 μM, the highest concentration tested, netarsudil resulted again in an IOP elevation by 3.65±0.86 mmHg (p <0.001), as seen in eyes with an intact TM.

**Figure 2:**
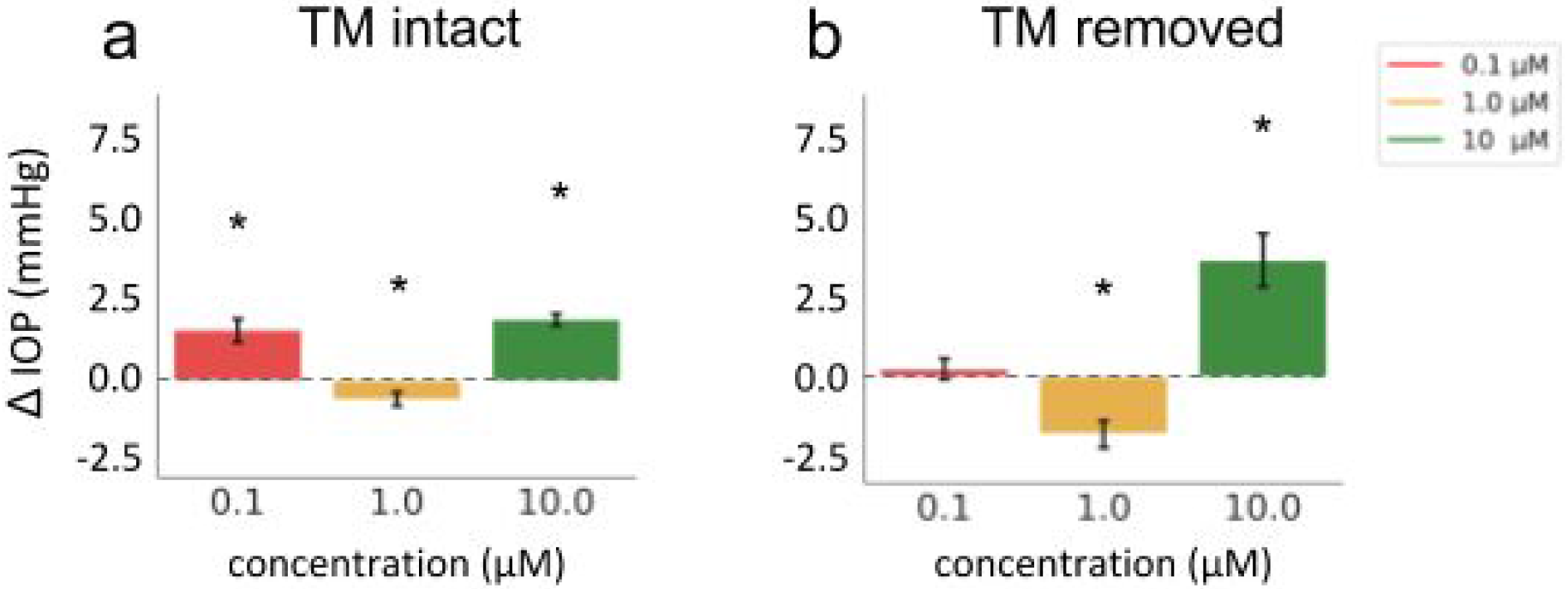
Netarsudil effect on IOP in a dose-dependent manner both with and without TM (* indicates significant difference from 0, one-sample t-test, P<0.05).

SD-OCT was able to measure CSA changes of vessels of the distal outflow tract (**Figure 3**). Corresponding to the IOP data, 0.1 μM netarsudil caused a 50±31% reduction of the CSA of perilimbal outflow tract vessels (**Figure 4**). In contrast, at 1 μM netarsudil, there was a 37±14% increase in CSA due to the dilation of outflow tract vessels. At 10 μM netarsudil, a constriction occurred again with a reduction of CSA by 43±7%.

**Figure 3:**
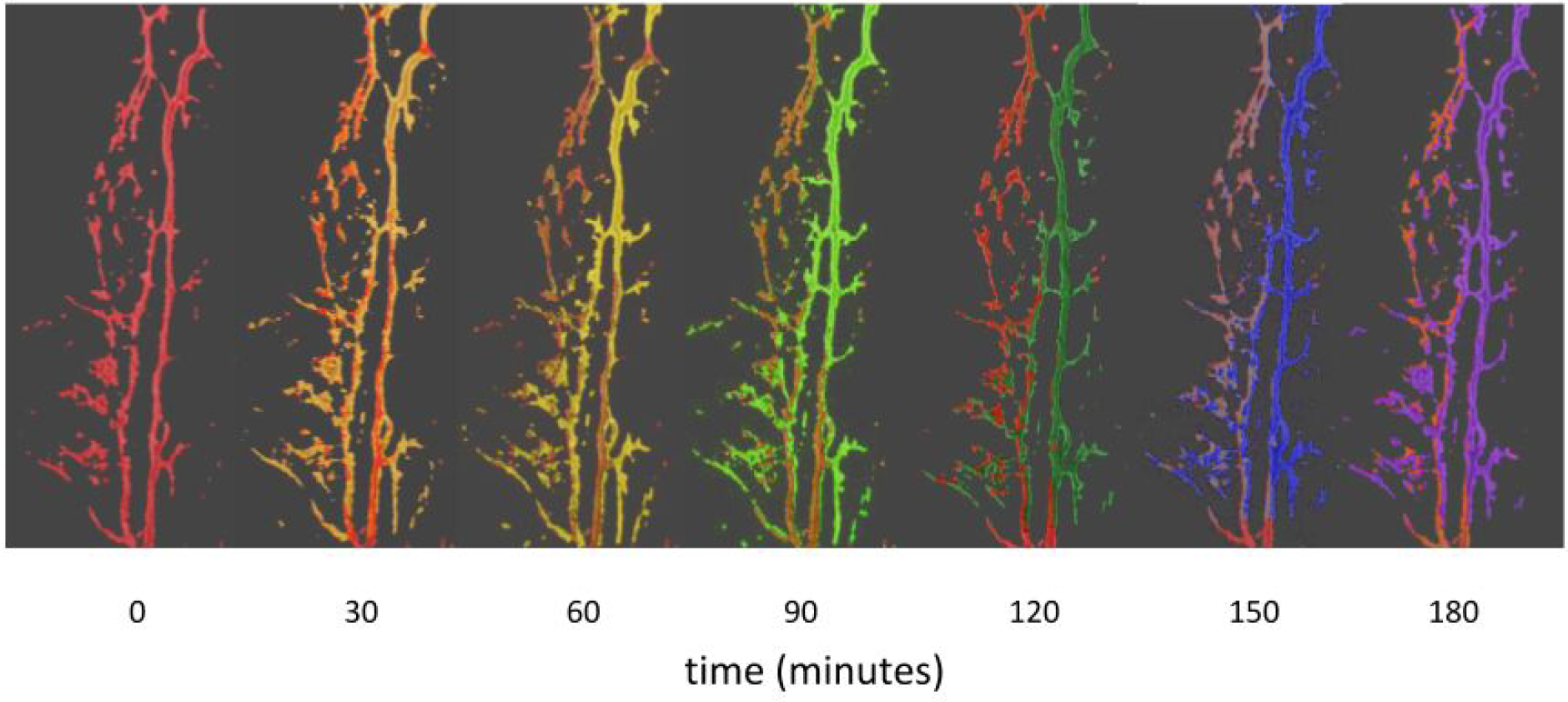
Overlay of SDOCT Amira snapshots of perilimbal outflow tract structures with a progressive dilation using an example at 1 μM netarsudil (red: 0 minutes, purple: 180 minutes). Overlay with color other than red indicates an increased vessel diameter.

**Figure 4:**
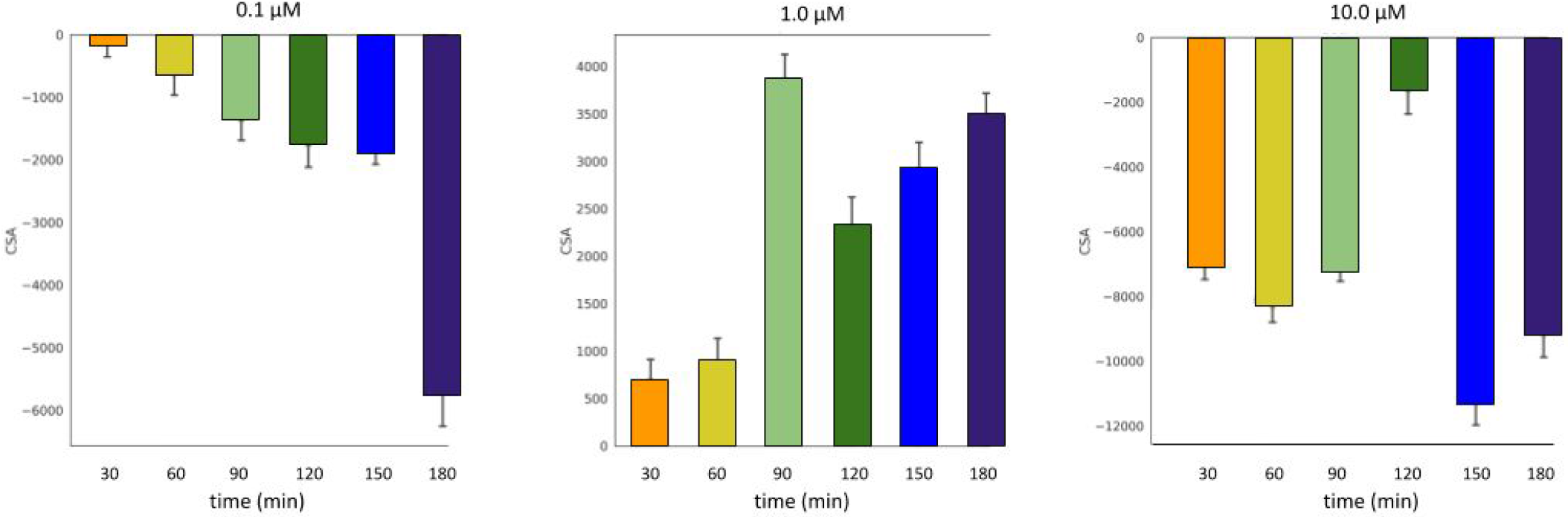
Change of cross section areas of outflow tract vessels over time at 0.1, 1.0 and 10.0 μM Netarsudil as measured by SDOCT (mean ± SD).

## Discussion

Recent evidence from clinical [20–23] and laboratory [10, 17] TM ablation studies demonstrated a significant post-TM outflow resistance that might be caused by a downstream regulatory mechanism [10, 17]. Only 0.3% of patients undergoing trabecular ablation in AIT achieve the predicted decrement in IOP to 8 mmHg akin to the level of episcleral veins [23]. An IOP glaucoma surgery calculator we derived from those data predicts glaucoma patients cannot achieve an IOP <18.6 mmHg without medications [24]. Even with topical glaucoma medications added back, TM ablation has been observed to have a failure rate of 28% within 12 months [20] for low IOP targets in moderate glaucoma while a higher preoperative IOP is correlated to an increased postoperative IOP.

In this study, we investigated the effect of netarsudil on the structure and function of the distal outflow tract at different concentrations. Pharmacological management of post trabecular outflow resistance holds promise to patients who fail microincisional angle surgery in glaucoma because rebounding of an initially low IOP or not achieving it to start with. First, we found that while a standard concentration of 1 μM of the Rho-kinase inhibitor netarsudil caused an IOP reduction and outflow vessel dilation, a lower (0.1 μM) and a higher concentration (10 μM) of netarsudil had the opposite effect, resulting in IOP elevation and outflow vessel constriction. The second finding of this study was that this effect did not require the TM but appeared to be mediated by distal outflow tract vessels.

The biochemistry and pharmacokinetics of netarsudil have been examined in animals [25, 26], and human eye models [15, 27] before recently entering clinical trials as a hypotensive agent for glaucoma [28, 29]. Netarsudil lowers IOP through a combination of three mechanisms, reduction of aqueous humor, increased trabecular facility and decreased episcleral venous pressure [28]. Aqueous humor production is reduced primarily through its action on the norepinephrine transporter, while the inhibition of Rho-kinase reduces stiffness [26] and stress fibers in the TM as seen with other Rho-kinase inhibitors [30]. The impact on vessel diameters is variable, but in clinical use, conjunctival hyperemia is a common observation [31] and probably caused by rendering vascular smooth muscle cells less sensitive to intracellular Ca^2+^ [32].

Our findings in porcine eyes are similar to Li et al.’s results in mouse [25] and Kiel et al.’s in rabbit eyes [33] who also observed dilation of outflow tract vessels with a corresponding pressure reduction. In preclinical studies, Rho-kinase inhibitors are potent inhibitors of ocular vasoconstriction [34], severe occlusive pulmonary arterial hypertension [35], and renal vasoconstriction [36] that hold potential for other chronic diseases. Release of a pathological post-trabecular outflow resistance might fit those if an outflow tract constriction and dysfunction can be confirmed in glaucoma. The dose-dependent IOP elevation and vasoconstriction by netarsudil found here suggest opposing rather than synergistic mechanisms and hints at a therapeutic window that might be more limited.

This ex vivo study had several limitations. The observed changes were statistically highly significant but small due to a perfusion rate that was only slightly higher than normal. We chose this rate out of concern to trigger NO release by excessive endothelial shear force [37–39]. In vivo, netarsudil might be converted more effectively into netarsudil-M1 than in this ex vivo model. Netarsudil-M1 is the esterase metabolite of netarsudil and has a greater potency [26]. Although the role of distal outflow tract resistance has been demonstrated in both porcine [10, 17] and human eyes [10], there might be species-dependent effect differences, especially as it relates to the extensive crosstalk between nitric oxide and RhoA/ROCK-signalling [40].

In conclusion, we found that netarsudil was able not only to decrease but also increase IOP in the porcine anterior segment culture depending on the concentration tested. The IOP changes matched the diameter changes of distal outflow tract vessels. The hyper- and hypotensive property of netarsudil persisted after TM removal.

